# Phosphorylation of the Anaphase Promoting Complex activator CDH1/FZR regulates the transition from Meiosis I to Meiosis II in mouse male germ cell

**DOI:** 10.1101/2020.03.13.990127

**Authors:** Nobuhiro Tanno, Shinji Kuninaka, Sayoko Fujimura, Kaho Okamura, Kazumasa Takemoto, Kimi Araki, Masatake Araki, Hideyuki Saya, Kei-ichiro Ishiguro

## Abstract

CDH1/FZR is an activator of Anaphase promoting complex/Cyclosome (APC/C), best known for its role as E3 ubiquitin ligase that drives the cell cycle. APC/C activity is regulated by CDK-mediated phosphorylation of CDH1 during mitotic cell cycle. Although the critical role of CDH1 phosphorylation has been shown mainly in yeast and *in vitro* cell culture studies, its biological significance in mammalian tissues *in vivo* remained elusive. Here, we examined the *in vivo* role of CDH1 phosphorylation using a mouse model, in which non-phosphorylatable substitutions were introduced in the putative CDK-phosphorylation sites of CDH1. Although ablation of CDH1 phosphorylation did not show substantial consequences in mouse somatic tissues, it led to severe testicular defects resulting in male infertility. In the absence of CDH1 phosphorylation, male juvenile germ cells entered meiosis normally but skipped meiosis II producing diploid spermatid-like cells. In aged testis, male germ cells were overall abolished, showing Sertoli cell-only phenotype. The present study demonstrated that phosphorylation of CDH1 is required for temporal regulation of APC/C activity at the transition from meiosis I to meiosis II, and for spermatoginial stem cell maintenance, which raised an insight into the sexual dimorphism of CDH1-regulation in germ cells.

## Introduction

Anaphase promoting complex/Cyclosome (APC/C) controls timely transitions of mitotic cell cycle phases by promoting ubiquitylation and degradation of many key cell cycle regulators (Acquaviva and Pines, 2006). APC/C activity is regulated by either of two co-activators CDC20 and CDH1/FZR, which determine the substrate specificity of ubiquitylation for each cell cycle phase (Peters, 2006). APC/C^CDC20^ activity appears in metaphase-to-anaphase transition, when it has an essential function in promoting chromosome segregation by mediating cyclin B1 and securin degradation. APC^CDH1^ is thought to regulate a wide range of cell cycle events, in which more than 30 proteins have been identified as APC/C^CDH1^ substrates (Ramanujan and Tiwari, 2016).

While CDC20 plays a role in APC/C activity at metaphase, when the cyclin-dependent kinase 1 (CDK1) activity is high, CDH1 contributes to APC/C activity when CDK1 activity is sustained at a low level (Peters, 2006). APC/C^CDH1^ activity is negatively regulated by CDK-mediated phosphorylation of CDH1 during mitotic cell cycle. At mitotic exit, reduction of CDK1 activity leads to phosphatase-mediated dephosphorylation of CDH1 which then binds and activates APC/C until late G1 phase (Wurzenberger and Gerlich, 2011). In yeast mitotic cell cycle, CDH1 is phosphorylated by increased level of CDK activity after late G1/S onward, which subsequently leads to dissociation of CDH1 from APC/C and its inactivation (Zachariae et al., 1998) (Jaspersen et al., 1999) (Blanco et al., 2000) (Robbins and Cross, 2010) (Hockner et al., 2016) (Ondracka et al., 2016). In mammals, HeLa cells transiently transfected with mutant CDH1 that lacked potential CDK1 phosphorylation sites, resulted in premature reduction of cyclin A and cyclin B with decrease in G2/M phase-cells (Kramer et al., 2000). Thus, CDK-mediated phosphorylation of CDH1 plays a crucial role in temporal regulation of APC/C activity during mitotic cell cycle. However, the physiological significance of CDH1 phosphorylation is yet to be examined in vertebrate mitotic tissues *in vivo*.

The meiotic cell cycle consists of one round of DNA replication followed by two rounds of chromosome segregation, producing haploid gametes from diploid cells. Although study in budding yeast elucidated that the role of APC/C in meiosis relies on the meiosis-specific co-activator AMA1, its homolog or counterpart does not exist in mammals. Instead, CDC20 and CDH1 contribute to the regulation of APC/C activity in mammalian meiosis. In mice, it was demonstrated that loss of CDH1 led to abnormal spermatogonial proliferation and defects in progression of meiotic prophase I (Holt et al., 2014).This indicates CDH1 is required for mammalian meiosis, but it remains elusive how APC/C activity regulated by phosphorylation of CDH1 is involved in meiotic cell cycle.

Here, we examined whether phosphorylation of CDH1 is required for the regulation of APC/C activity during mitotic and meiotic cell cycle *in vivo*, using knockin mice that carries non-phosphorylatable mutations of CDH1. Our study demonstrates that phosphorylation of CDH1 is required for spermatoginial stem cell maintenance for a long period of time. Furthermore, we show that CDH1 phosphorylation has a crucial role in the regulation of APC/C activity during meiosis I-meiosis II transition in juvenile male but not in female. Sexual dimorphism in the requirement of CDH1 phosphorylation raised an insight into different modes of meiosis I-to-meiosis II progression in spermatocytes and oocytes.

## Results

### Generation of *Cdh1*^9A/9A^ knockin mice

It has been shown that human CDH1 is phosphorylated during mitotic cell cycle. Ectopic expression of non-phosphorylatable CDH1 mutant in HeLa cells resulted in formation of constitutively active APC/C^CDH1^ and a reduction of G2/M phase (Kramer et al., 2000). To analyze the physiological role of CDH1-phosphorylation, we generated a mouse model, *Cdh1*^9A/9A^ knockin (*Cdh1*^9A/9A^ KI) mice, in which the nine putative CDK-phosphorylation sites of Ser/Thr residues in CDH1 were substituted with alanine amino acids (Fig.1A-C). *Cdh1*^9A/9A^ KI allele was generated by Cre-mediated site-specific recombination of full length cDNA encoding mutant *Cdh1*^9A^ into the endogenous *Cdh1* locus using an exchangeable gene-trap (GT) line (Taniwaki et al., 2005) (Naoe et al., 2010). For the control mouse line, full length cDNA cassette encoding wild type (WT) *Cdh1* was inserted into the targeted locus in the same manner, generating *Cdh1*^Gt wt/Gt wt^ KI mice (Fig.1A). To distinguish *Cdh1*^Gt wt^ KI allele from natural WT *Cdh1* allele, hereafter we refer to natural WT *Cdh1* allele as *Cdh1*^+^. The expression levels of CDH1-Gt wt and CDH1-9A proteins in the corresponding KI testes were comparable to the CDH1 expression level in natural WT *Cdh1*^+/+^ testes at postnatal day18 (P18) (Fig.1D). Further, CDC20, APC/C subunits (CDC27/APC3) and the canonical substrates of APC/C (Cyclin B1, PLK1) overall showed similar expression levels in natural WT *Cdh1*^+/+^, *Cdh1*^9A/9A^ KI and *Cdh1* ^Gt wt/Gt wt^ KI testes (Fig.1D). We also confirmed *Cdh1*^Gt wt/Gt wt^ KI mice showed normal fertility as natural WT *Cdh1*^+/+^, indicating *Cdh1*^Gt wt^ KI allele functions in a manner indistinguishable from natural WT *Cdh1*^+^ allele. Immunoprecipitation of CDC27/APC3, a core subunit of APC/C, demonstrated that the non-phoshorylatable CDH1-9A and control CDH1-Gt wt proteins expressed from the corresponding KI alleles were indeed incorporated in APC/C of *Cdh1*^9A/9A^ and *Cdh1*^Gt wt/Gt wt^ KI testes, respectively (Fig.1E). We noticed that the level of CDC20 incorporated in APC/C in *Cdh1*^9A/9A^ testis was less than that in natural WT *Cdh1*^+/+^ and *Cdh1*^Gt wt/Gt wt^ KI testes. This implies that persistent inclusion of CDH1-9A in APC/C impedes the association of CDC20 to APC/C.

**Figure 1.**
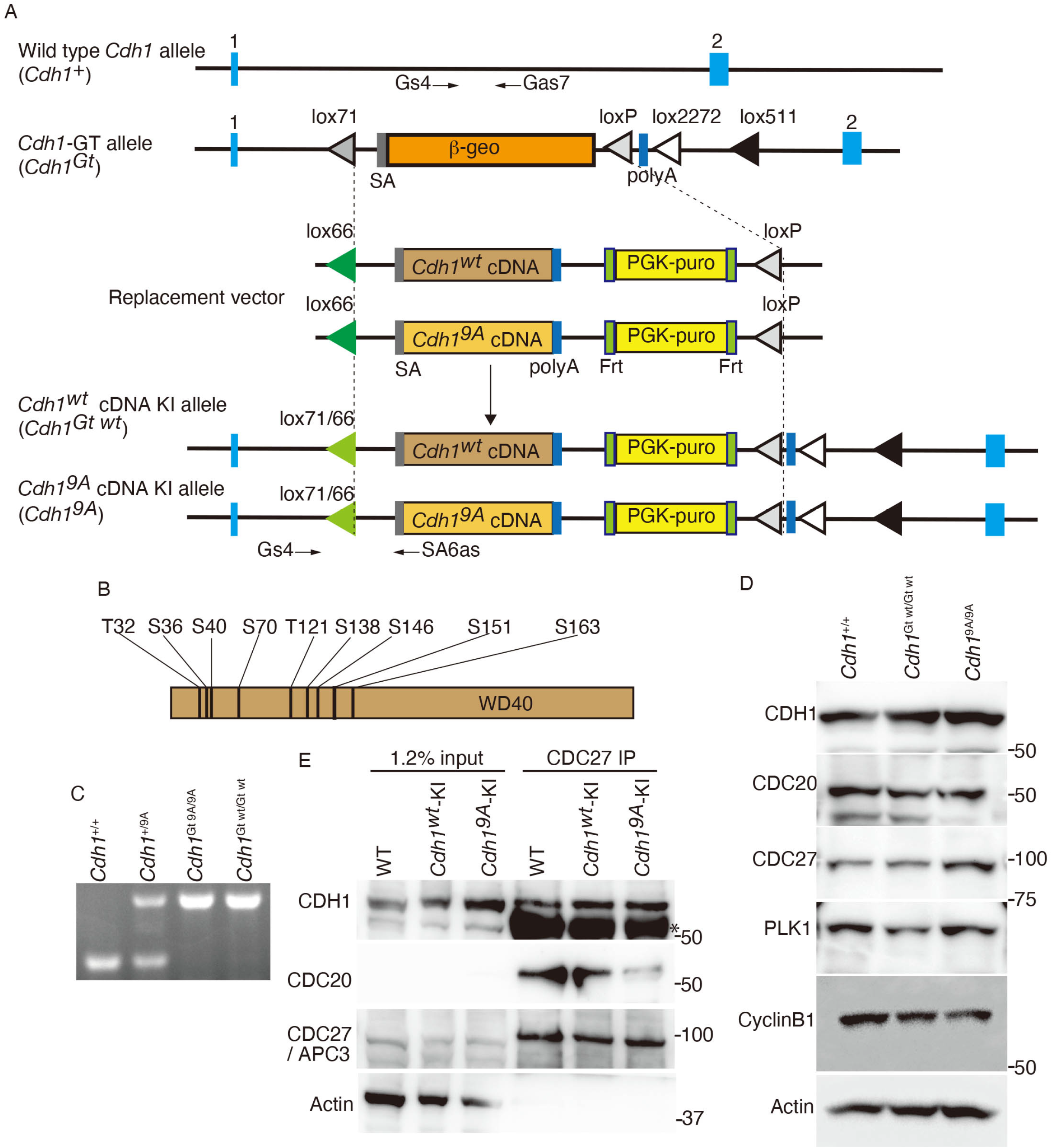
Generation of *Cdh1*^9A/9A^ knockin mice. **(A)**Schematic representation of the WT *Cdh1*^+^ allele, the Gene trapped (GT) allele *Cdh1*^GT^, the knock-in (KI) alleles of *Cdh1*^GT wt^ and *Cdh1*^9A^ and the replacement vectors possessing *Cdh1*^GT wt^ or *Cdh1*^9A^ cDNA. *Cdh1* ^GT^ allele was generated by integration of the exchangeable Gene Trap (GT) vector in the 1st intron of the endogenous *Cdh1* locus (Naoe et al., 2010). Cre-mediated recombination between *Cdh1*^GT^ allele and the replacement vector generated *Cdh1*^9A^ KI allele. The control *Cdh1*^wt^ allele was generated between *Cdh1*^GT^ allele and the replacement vector possessing *Cdh1* ^wt^ cDNA in the same manner. SA: splicing acceptor, Frt: Flippase recombination site, polyA: poly adenylation signal, lox P and its variant recombination sites are indicated by triangles. Light blue rectangles indicate the exons of *Cdh1* locus. Arrows indicate the PCR primers for genotyping. (**B**) Nine putative CDK-phosphorylated sites of Ser and Thr residues in CDH1, where Ala substitutions were introduced. (**C**) PCR genotyping of genomic DNA from *Cdh1*^+/+^, *Cdh1*^+/9A^, *Cdh1*^9A/9A^ and *Cdh1*^Gt wt/Gt wt^ KI mice. **(D)** Western blot of testis extracts from *Cdh1*^+/+^, *Cdh1*^Gt wt/Gt wt^ and *Cdh1*^9A/9A^ testes (P18), probed by antibodies as indicated. **(E)** Western blot of immunoprecipitates from testis extracts of *Cdh1*^+/+^, *Cdh1*^Gt wt/Gt wt^ and *Cdh1*^9A/9A^ mice using anti-CDC27 antibody. * indicates nonspecific bands cross-reacted with IgG. See also Fig S3 for the uncropped images.

### Introduction of non-phosphorylatable mutations of CDH1 leads to male infertility

Inter-crossing of heterozygous (*Cdh1*^9A/+^) males and females produced average 7.285 litter size with normal number of litters of *Cdh1*^+/+^, *Cdh1*^+/9A^ and *Cdh1*^9A/9A^ genotypes expected by mendelian ratio (total *Cdh1*^+/+^:11 pups, *Cdh1*^+/9A^: 28 pups and *Cdh1*^9A/9A^: 12 pups from 7 pairs of intercrossing. *p* =0.9557 by chi-square test). The homozygous *Cdh1*^9A/9A^ KI mice developed normally without any overt phenotype in adult somatic tissues, suggesting that CDH1-9A showed little impact on mouse development and mitotic cell cycle of mouse somatic tissues.

Notably, although testes in juvenile *Cdh1*^9A/9A^ KI males did not show apparent difference from those of control WT *Cdh1*^+/+^ or *Cdh1*^+/9A^, defects in male reproductive organs were evident with smaller-than-normal testes at adulthood (Fig. 2A).

**Figure 2.**
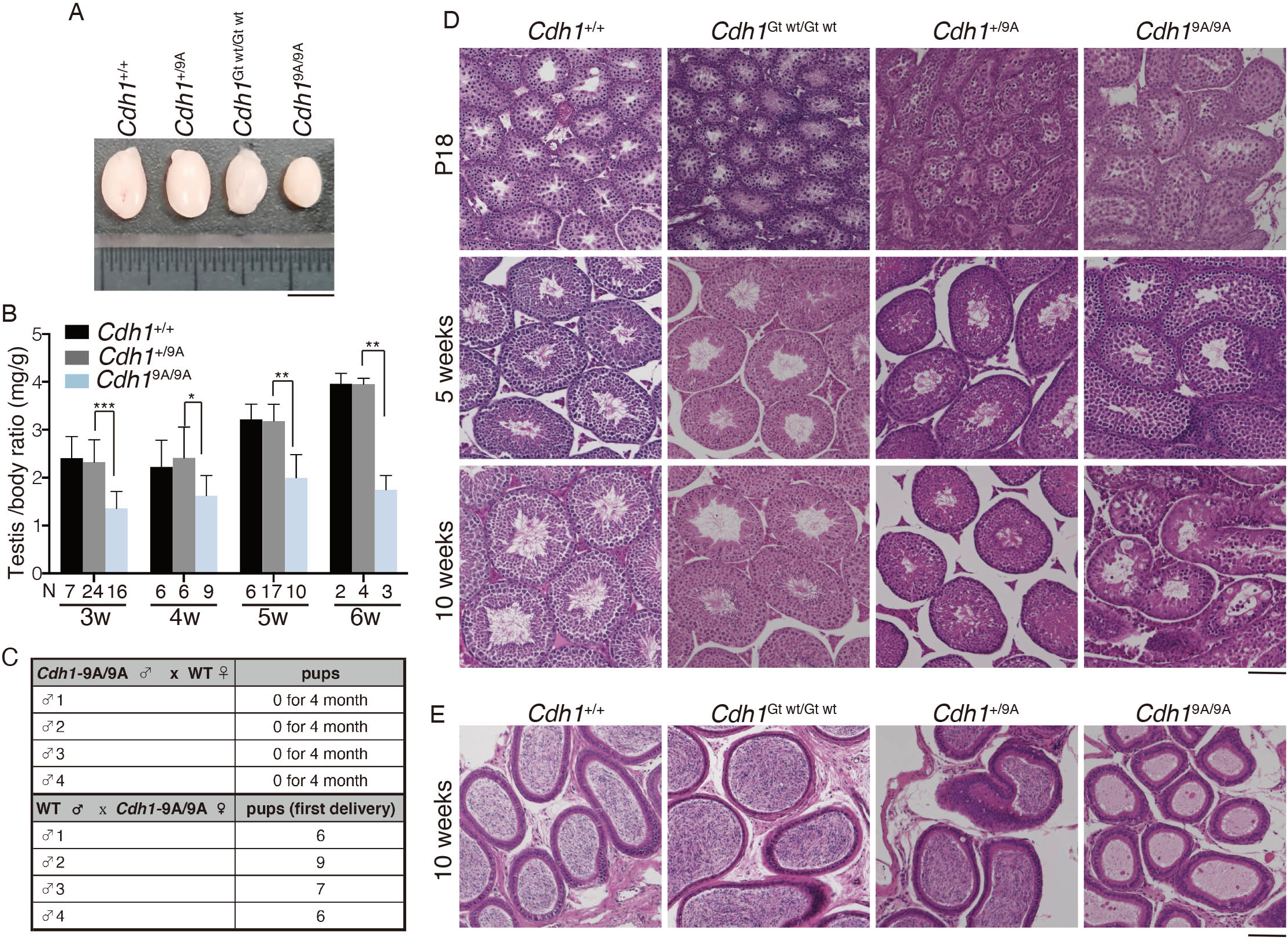
Spermatogenesis is defective in *Cdh1*^9A/9A^ knockin mice. **(A)** Gross morphology of representative testes from 8-week-old control *Cdh1*^+/+^, *Cdh1*^+/9A^, *Cdh1*^Gt wt/Gt wt^ and *Cdh1*^9A/9A^ KI mice. Scale bar, 5mm. **(B)** Testis /body weight ratio (mg/g) of control *Cdh1*^+/+^, *Cdh1*^+/9A^ and *Cdh1*^9A/9A^ KI testes at the indicated ages. N: number of animals examined. *p*-values are shown (paired t-test). ***: *p* <0.001, **: *p* <0.01, *: *p* <0.05 **(C)** Number of pups born by mating *Cdh1*^9A/9A^ KI males with wild type females, and *Cdh1*^9A/9A^ KI females with wild type males to examine fertility. **(D)** Hematoxylin and Eosin stained histological sections of seminiferous tubules in WT, *Cdh1*^Gt wt/Gt wt^, *Cdh1*^+/9A^ and *Cdh1*^9A/9A^ KI mice at the indicated ages. **(E)** Hematoxylin and Eosin stained sections of the epididymis in WT, *Cdh1*^Gt wt/Gt wt^, *Cdh1*^+/9A^ and *Cdh1*^9A/9A^ KI mice at the age of 10 weeks. Scale bars: 100 μm except A.

Remarkably, testes in the *Cdh1*^9A/9A^ KI males at 3 week or older were significantly smaller compared to the age-matched *Cdh1*^+/+^ males (Fig. 2B). Since it is known that the first wave of spermatogenesis completes meiotic prophase and reaches to meiotic divisions generating haploid spermatids at earliest P18, the observation in *Cdh1*^9A/9A^ KI males suggests that the primary defect of *Cdh1*^9A/9A^ KI males emerges during or immediately after meiotic prophase.

To determine whether the introduction of non-phosphorylatable mutations in CDH1 affected male fertility, 8-week-old *Cdh1*^9A/9A^ KI males, control WT *Cdh1*^+/+^ males and *Cdh1*^Gt wt/Gt wt^ KI males were mated with C57BL6 wild-type females over a period of 4 months. While the control WT *Cdh1*^+/+^ and *Cdh1*^Gt wt/Gt wt^ KI males produced normal size of litters over this period, the *Cdh1*^9A/9A^ KI males showed infertility over a period of 4 months (Fig. 2C). Since the control *Cdh1*^Gt wt/Gt wt^ KI mice showed normal fertility, it is tenable that the infertility in the *Cdh1*^9A/9A^ KI males was attributed to the introduction of non-phosphorylatable mutations in CDH1. Since heterozygous *Cdh1*^9A/+^ KI males showed normal fertility, the phenotype exhibited by the *Cdh1*^9A^ KI allele was recessive rather than dominant negative in terms of male fertility (Fig. 2B). In contrast to the male phenotype, *Cdh1*^9A/9A^ KI females showed normal fertility when mated with C57BL6 wild-type males (Fig. 2C) and no overt defects in adult ovaries (Fig. S1). Thus the infertility caused by non-phosphorylatable mutations in CDH1 was male specific. Histological analysis revealed that obvious difference was not observed between *Cdh1*^9A/9A^ KI and the control seminiferous tubules at P18 (Fig. 2D), suggesting that the first wave of spermatogenesis normally progressed through meiotic prophase in *Cdh1*^9A/9A^ KI testis. However, at 5 week-old onward, spermatozoa were absent in the *Cdh1*^9A/9A^ KI seminiferous tubules, whereas spermatogonia, spermatocytes and round spermatid-like cells appeared. This observation was in sharp contrast to those of the age-matched natural WT *Cdh1*^+/+^ and the control *Cdh1*^Gt wt/Gt wt^ KI males (Fig. 2D). Thus, spermatogenesis was severely impaired in the *Cdh1*^9A/9A^ KI seminiferous tubules. Accordingly, spermatozoa were completely absent in *Cdh1*^9A/9A^ KI epididymis (Fig.2E), corroborating the infertility of the *Cdh1*^9A/9A^ KI males. These observations suggested that although the germ cells had undergone spermatogonial differentiation and meiosis, they failed to progress spermiogenesis beyond round spermatids in the *Cdh1*^9A/9A^ KI seminiferous tubules at younger age. Therefore, we reasoned that spermatogenesis defects in *Cdh1*^9A/9A^ KI male could be attributed to the failure of differentiation in round spermatids or in later stage during the juvenile age. Altogether, phosphorylation of CDH1 is required for normal progression of spermatogenesis.

### Phosphorylation of CDH1 is required for meiosis I-meiosis II transition in male

Given that spermatogenesis defects in *Cdh1*^9A/9A^ KI male might arise during or after meiosis in the juvenile age, we further investigated at which stage the primary defect appeared in juvenile *Cdh1*^9A/9A^ KI. We analyzed the progression of spermatogenesis in juvenile *Cdh1*^9A/9A^ KI males by immunostaining using stage specific markers. Consistent with above observations, immunostaining analysis with antibodies against SYCP3 (a component of meiotic chromosome axis) along with γH2AX (a marker of double strand break for meiotic recombination) or SYCP1 (a marker of homologous chromosome synapsis) demonstrated that spermatocytes underwent meiotic recombination and homologous chromosome synapsis in juvenile *Cdh1*^9A/9A^ KI males, which were comparable to age-matched WT (Fig. S2A, B). The *Cdh1*^9A/9A^ KI spermatocytes reached mid to late pachytene stage as indicated by testis-specific histone H1t, which were also comparable to age-matched WT (Fig. S2C). Thus, spermatocytes progressed the meiotic prophase normally in juvenile *Cdh1*^9A/9A^ KI males. We further investigated the meiotic processes before and after the first meiotic division. Spread nuclei from the testes were immunostained with antibodies against SYCP3 along with MEIKIN, a kinetochore marker of late pachytene-metaphase I (Kim et al., 2015). In *Cdh1*^9A/9A^ KI, spermatogenesis reached metaphase I (centromeric SYCP3+/MEIKIN+) and interkinesis (bar-like SYCP3+/MEIKIN−), an interphase-like stage between meiosis I and meiosis II (Parra et al., 2004), (Fig. 3A) suggesting that at least meiosis I was complete in *Cdh1*^9A/9A^ KI spermatocytes. Intriguingly, although *Cdh1*^9A/9A^ KI spermatocytes reached interkinesis at the most advanced stage, we noticed that atypical round spermatid-like cells appeared in the *Cdh1*^9A/9A^ KI (Fig. 3B). Whereas WT round spermatids mostly exhibited single chromocenters (the clusters of centromeric heterochromatin), *Cdh1*^9A/9A^ KI round spermatid-like cells showed multiple number of chromocenters with larger size of nuclei (Fig. 3C). We also observed a small fraction of elongated spermatid-like cells with aberrant shape in *Cdh1*^9A/9A^ KI testis (Fig. 3D). Notably, since *Cdh1*^9A/9A^ KI spermatid-like cells were PNA lectin (a marker of spermatid) positive, spermiogenesis program at least in part progressed in *Cdh1*^9A/9A^ KI atypical spermatid-like cells. Crucially, flow cytometry analysis of testicular cells from 8-week-old *Cdh1*^9A/9A^ KI testes demonstrated complete absence of the haploid (1N) population, despite the existence of spermatid-like cells (Fig. 3E). Therefore, we reasoned that the observed primary defect in *Cdh1*^9A/9A^ KI spermatogenesis derived from a process during meiosis I-meiosis II transition rather than a failure in post-meiotic differentiation into elongated spermatid. Because we did not observe any population with 1N DNA content, round spermatid-like cells were diploid (2N) after normal meiosis I chromosome segregation. It is plausible that secondary spermatocytes failed to progress from the interkinesis stage to meiosis II in *Cdh1*^9A/9A^ KI testes. Furthermore, a subpopulation of those round spermatid-like cells was TUNEL positive in *Cdh1*^9A/9A^ KI seminiferous tubules, suggesting that aberrant round and elongated spermatid-like cells were eliminated at least in part by apoptosis (Fig. 3F).

**Figure 3.**
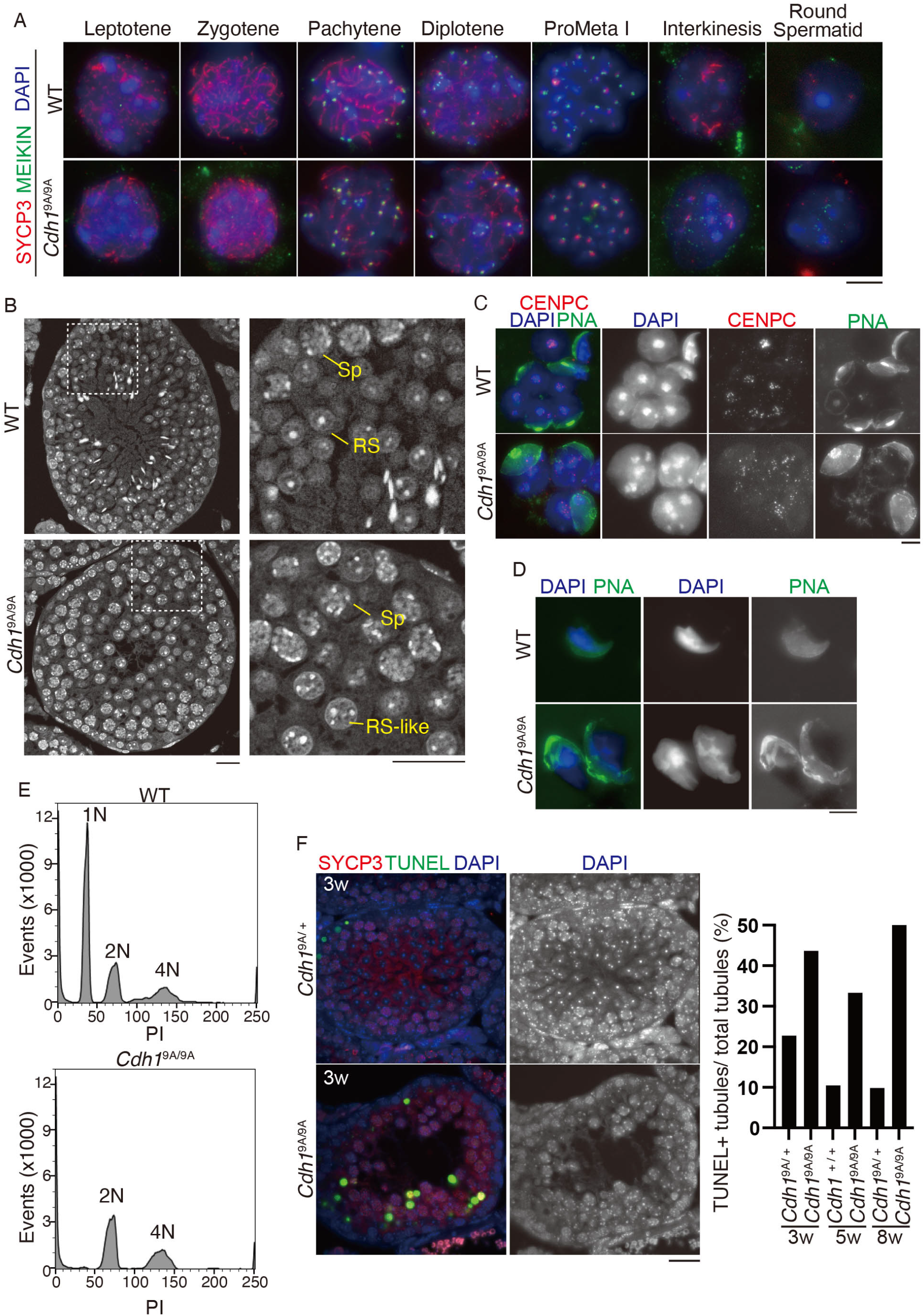
*Cdh1*^9A/9A^ knockin testes show defect in meiosis I-meiosis II transition. **(A)** Spread nuclei from the testes of wild type and *Cdh1*^9A/9A^ KI mice were immunostained for SYCP3, MEIKIN, and DAPI. Scale bar: 5 μm. **(B)** Seminiferous tubule sections in *Cdh1*^+/9A^ and *Cdh1*^9A/9A^ KI mice (6-week-old) were DAPI-stained. Enlarged images are shown on the right to highlight the round-spermatid like cells in *Cdh1*^9A/9A^ KI mice. Sp: spermatocyte, RS: round spermatid, RS-like: round spermatid-like cell. Scale bar: 20 μm. **(C)** The round spermatid and the round spermatid-like cells from the testes of WT and *Cdh1*^9A/9A^ KI mice, respectively, were immunostained for Centromere protein-C (CENPC), PNA lectin and DAPI. Scale bar: 5 μm. **(D)** The elongated spermatid and the elongated spermatid-like cells from the testes of WT and *Cdh1*^9A/9A^ KI mice, respectively, were immunostained for PNA lectin and DAPI. Scale bar: 5 μm. **(E)** Flow cytometry histogram of propidium-iodide-stained testicular cells isolated from *Cdh1*^+/9A^ and *Cdh1*^9A/9A^ KI mice (P28). N indicates DNA content in the testicular cell population. **(F)** Seminiferous tubule sections from *Cdh1*^+/9A^ and *Cdh1*^9A/9A^ KI mice (3 week old) were stained for SYCP3, TUNEL and DAPI. Quantification of the seminiferous tubules that have TUNEL+ cells per total tubules in control (WT or *Cdh1*^+/9A^) and *Cdh1*^9A/9A^ KI testes (3w, 5w, 8w: each n = 1). Scale bar: 25 μm.

### Phosphorylation of CDH1 is required for the entry into meiosis II in male meiosis

Given that *Cdh1*^9A/9A^ KI spermatocytes undergo meiosis I but do not produce haploid cells, the round spermatid-like cells may be a consequence of the failure of meiosis II. Homologous chromosomes are segregated in meiosis I, whereas sister chromatids are separated in meiosis II. In order to examine the chromosome composition in *Cdh1*^9A/9A^ KI round spermatid-like cells, we performed immuno-FISH assays using a probe that detects a specific DNA sequence in the mid-arm region of chromosomes 3. In both WT and *Cdh1*^9A/9A^ KI mice, interkinesis spermatocytes showed a pair of FISH signals, indicating that homologous chromosomes were disjoined after meiosis I (Fig. 4A). In contrast, while WT round spermatids showed a single FISH signal, most of *Cdh1*^9A/9A^ KI round spermatid-like cells showed a pair of FISH signals (Fig. 4B), suggesting that in *Cdh1*^9A/9A^ KI interkinesis spermatocytes failed to enter or complete meiosis II, and consequently led to non-disjunction of sister chromatids. Thus, these results suggested that the progression from interkinesis to meiosis II was regulated by phosphorylation of CDH1 in male meiosis. In mitotic cell cycle, constitutively active APC/C^Cdh1^ reduces cyclin A and cyclin B levels prematurely with decrease in G2/M phase cells (Kramer et al., 2000). Our observations in *Cdh1*^9A/9A^ KI male implies that without phosphorylation of CDH1, APC/C^Cdh1^ substrates might undergo premature degradation, which impairs male meiosis I – meiosis II transition.

**Figure 4.**
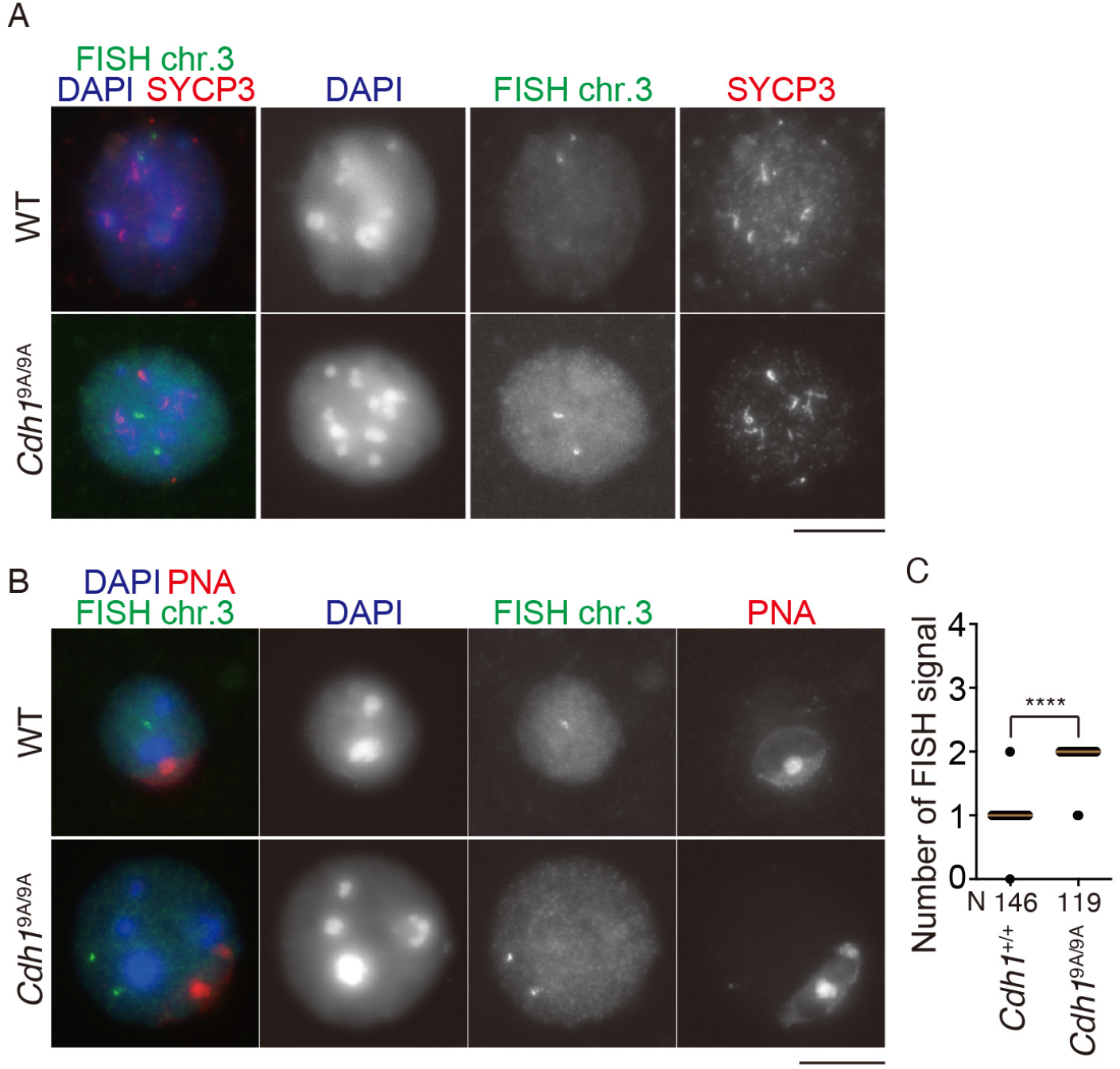
*Cdh1*^9A/9A^ KI spermatocytes lead to sister chromatid non-disjunction. **(A)** Interkinesis spermatocytes from wild type and *Cdh1*^9A/9A^ KI immunostained with SYCP3 (red) were subjected to FISH with a point probe (Chr.3) (green). **(B)** WT round spermatids and *Cdh1*^9A/9A^ KI round spermatid-like cells (14 week old) immunostained with SYCP3 and PNA were subjected to FISH with a point probe (Chr.3) (green). **(C)** The number of FISH signals in WT round spermatids and *Cdh1*^9A/9A^ KI round spermatid-like cells are represented in a scatter plot with medians. ****: *p* <0.0001 (t-test). N: number of examined cells. Scale bars: 5 μm.

### Phosphorylation of CDH1 is required for maintaining spermatogonial stem cell population

We noticed that the loss of germ cells was more severe in aged *Cdh1*^9A/9A^ KI seminiferous tubules, with residual spermatogonia, spermatocytes and Sertoli cells remaining along the basement compartment of the tubules (Fig.5A), which was accompanied by complete absence of spermatozoa in the epididymis (Fig.5B). Immunostaining of the aged *Cdh1*^9A/9A^ KI seminiferous tubules revealed that TRA98 positive cells were lost while SOX9 positive cells remained in a subpopulation of the tubules, suggesting that germ cell populations were depleted (Fig.5C). This trend was further augmented at older age in *Cdh1*^9A/9A^ KI seminiferous tubules (Fig.5D). The aged *Cdh1*^9A/9A^ KI seminiferous tubules showed a severe loss of male germ cells, which was similar to Sertoli cell-only phenotype. This implied spermatogonial stem cell population was depleted over a long period of time in *Cdh1*^9A/9A^ KI testis. Therefore, it is plausible that phosphorylation of CDH1 is required also for maintaining the spermatogonial stem cell population.

**Figure 5.**
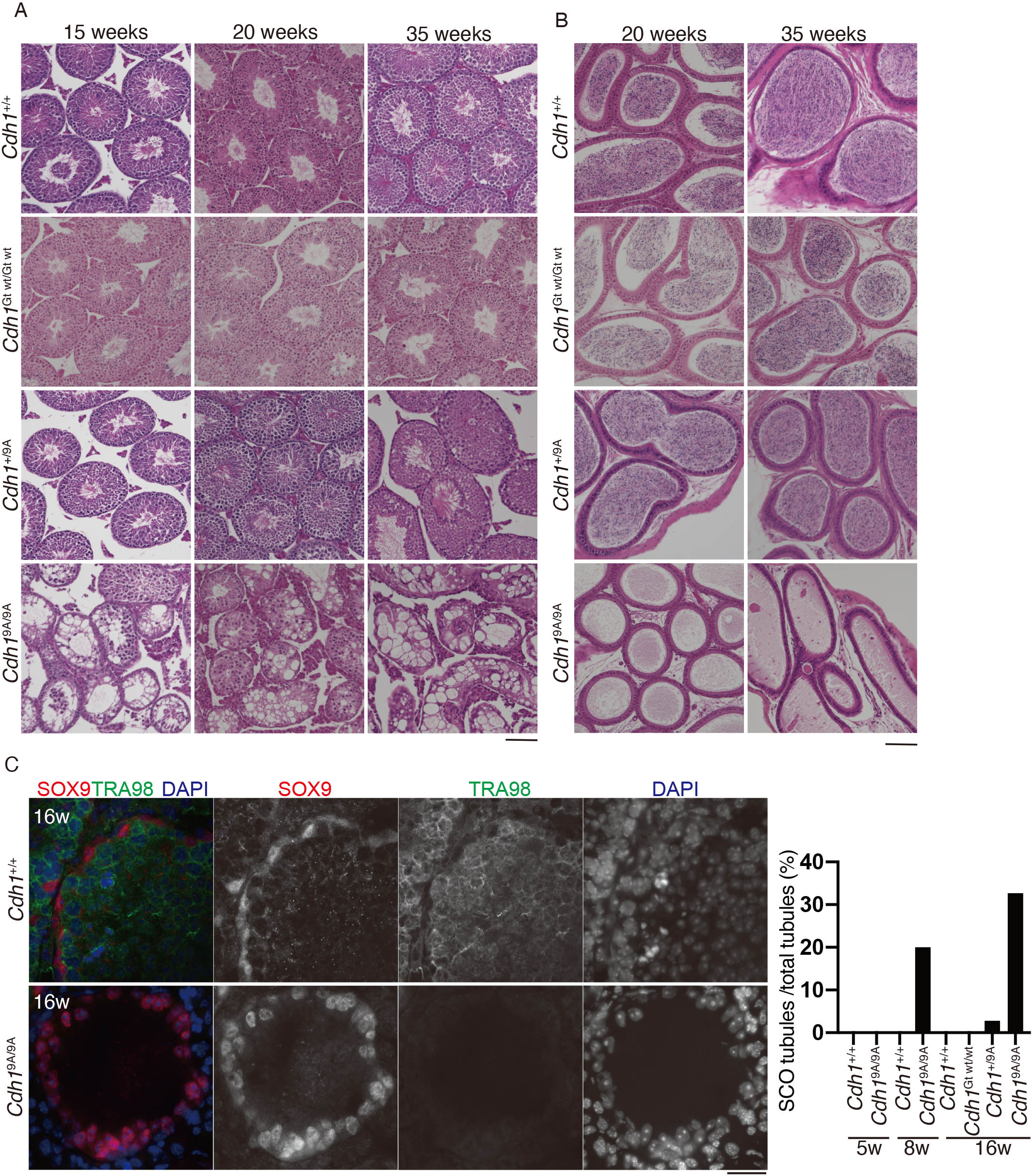
Sertoli cell-only phenotype appeared in aged *Cdh1*^9A/9A^ KI testes. **(A)** Hematoxylin and Eosin stained histological sections of seminiferous tubules in WT, *Cdh1*^Gt wt/Gt wt^, *Cdh1*^+/9A^ and *Cdh1*^9A/9A^ KI mice at the indicated ages. Scale bar: 100 μm. **(B)** Hematoxylin and Eosin stained sections of the epididymis in WT, *Cdh1*^Gt wt/Gt wt^, *Cdh1*^+/9A^ and *Cdh1*^9A/9A^ KI mice at the indicated ages. Scale bar: 100 μm. **(C)** Seminiferous tubule sections of indicated genotypes (5, 8 and 16-week-old) were stained for SOX9 (Sertoli cell marker), TRA98(germ cell marker) and DAPI. **(D)**Quantification of the seminiferous tubules that show Sertoli cell-only (SCO) phenotype per total tubules. Scale bar: 25 μm.

## Discussion

Contrary to previous observation in human cell culture study (Kramer et al., 2000), *Cdh1*^9A/9A^ KI mice showed no overt phenotype in somatic tissues. This suggests that somatic cells undergo normal mitotic cell cycle in *Cdh1*^9A/9A^ KI mice, despite the constitutive activity of APC/C^CDH1^. It is possible that premature degradation of APC/C^CDH1^ specific substrates, such as cyclin A, cyclin B, DNA replication regulators (CDC6 and Geminin) and mitosis regulators (Aurora A, Aurora B, PLK1, CDC20 and SGO1), might be compensated by robust balance of cell cycle regulators in somatic tissues in the mutant mice.

Crucially, we have shown that introduction of non-phosphorylatable mutations of CDH1 results in spermatogenesis defect and subsequent male infertility (Fig 2), which cannot be compensated as in mitotic cell cycle. The defects in the *Cdh1*^9A/9A^ KI testes primarily arose from a process during meiosis I-meiosis II transition producing diploid round spermatid-like cells (Figs 3 and 4). This phenotype is reminiscent of the observation that large round spermatid-like cells were accumulated in *Meikin* KO testes, where meiosis II was skipped as a result of failure in reductional chromosome segregation during meiosis I (Kim et al., 2015). The presence of diploid round spermatid-like cells suggested that the *Cdh1*^9A/9A^ KI spermatocytes were not simply arrested before meiosis II. At least, the *Cdh1*^9A/9A^ KI spermatocytes reached interkinesis but failed to enter or complete meiosis II. It should be mentioned that despite the aberrant cell cycle regulation at meiosis II, the *Cdh1*^9A/9A^ KI spermatocytes still showed an ability to undergo spermiogenesis. This implies that meiotic cell cycle program is genetically separable from spermiogenesis.

In contrast, non-phosphorylatable mutation of CDH1 exhibits apparently no impact on female fertility, showing sexual dimorphism of the *Cdh1*^9A/9A^ allele. This might be attributed to the sexual difference in the mode of meiosis I-meiosis II transition. In male, spermatocytes pass through a transient interphase-like state (Parra et al., 2004) with chromosome de-condensation and reassembly of nuclear membrane after completion of meiosis I. In female, chromosomes are persistently condensed without nuclear compartmentalization after completion of meiosis I until metaphase II. We hypothesize that this sexual difference is accounted for, at least in part, by CDK activity during meiosis I-meiosis II transition. While CDK activity remains high enough to prevent reassembly of nuclear membrane and chromosome decondensation after meiosis I until metaphase II in female, maybe it decreases once after meiosis I and then increases until metaphase II in male. Since CDK-mediated phosphorylation of CDH1 inactivates APC/C ^CDH1^ through preventing its association with APC/C (Zachariae et al., 1998) (Jaspersen et al., 1999) (Blanco et al., 2000) (Kramer et al., 2000), it is possible that constitutively active APC/C ^CDH1-9A^ compromise the levels of its critical substrates, such as cyclin A and cyclin B, which in turn prevent the CDK activity required for nuclear breakdown at interkinesis and entry into meiosis II. Alternatively, constitutive binding of CDH1-9A may compete with another activator CDC20 for the association with APC/C before meiosis II, which in turn raises a situation that APC/C ^CDC20^ specific substrates fail to undergo destruction in meiosis II.

We have also shown that the *Cdh1*^9A/9A^ KI testes show Sertoli-cell only phenotype at older age (Fig 5). Depletion of germ cells in *Cdh1*^9A/9A^ KI testes may imply that constitutive APC/C ^Cdh1-9A^ activity enforces persistent transition of spermatogonial stem cell population into G1/S, preventing their entry into G0 quiescence state. Thus, CDK-mediated phosphorylation of CDH1 is required for sustaining spermatogonial stem cells over a long period of time.

Overall, temporal regulation of CDH1 by phosphorylation is essential for the maintenance of spermatogonial stem cell population and meiosis II transition in male germ cells.

## Acknowledgments

The authors thank Tomohiko Wakayama, Yuki Takada (Kumamoto University) for technical support and valuable advice. This work was supported in part by KAKENHI grant #16H01257, #16H01221, #17H03634, #18K19304, #19H05245, #19H05743, #JP16H06276 from MEXT, Japan (to K.I.), the IMEG program of the Joint Usage/Research Center for Developmental Medicine (to H.S.).

## Author contributions

N.T. and K.I. performed the experiments. K.A. and M.A. designed the knockin mice and supported the mice experiments. S.F. performed histological analyses. K.T., K.O. supported the experiments. H.S., S.K. and K.I. supervised experiments and conducted the study. K.I. and N.T. wrote the manuscript.

## Materials & Methods

### Animal experiments

*Cdh1*^9A/9A^ KI and *Cdh1*^wt/wt^ mice were 129/C57BL6 mixed genetic background. Whenever possible, each knockin animal was compared to littermates or age-matched non-littermates from the same colony, unless otherwise described. Since *Cdh1*^+/+^, *Cdh1*^Gt wt/Gt wt^ KI, *Cdh1*^+/9A^ KI mice showed indistinguishable phenotype in fertility and testes morphology, they were used as controls when compared to *Cdh1*^9A/9A^ KI mice. Animal experiments were approved by the Institutional Animal Ethics Committees of Keio University (approval 16037-1) and Kumamoto University (approval F28-078, A2020-006, A30-001, A28-026).

### Generation of *Chd1* knock-in mouse and genotyping

*Cdh1*^+/GT^ allele was generated in TT2 embryonic stem (ES) cell line (Yagi et al., 1993) by integration of the pU-17 exchangeable GT vector (Taniwaki et al., 2005) into the *Cdh1/Fzr1* locus (Naoe et al., 2010). Characterization of the vector insertion site was performed by 5’ rapid amplification of cDNA ends (5’ RACE) and plasmid rescue experiments. Genotyping of the mutant mice was performed using a PCR protocol based on the primers Gs4 (5’ -CCTCCACTACAGCAGCACG-3’), Gas7 (5’-CTCCAAGGCCTTTGTGAGGC-3’), and SA6as (5’-CCGGCTAAAACTTGAGACCTTC-3’). For detection of the *Cdh1*–*β-geo* fusion mRNA, oligo(dT)-primed cDNAs derived from mutant mice were subjected to PCR using the primers 5NC-s (5’-TGTTCCTGGGACCGGCGGGAAC-3’) and LZUS-3 (5’-CGCATCGTAACCGTGCATCT-3’). The amplification product was cloned into the TA cloning vector and sequenced.

To produce ES cells in which the *β-geo* gene cassette of *Cdh1*^+/GT^ cells was replaced with cDNA encoding mouse wild type *Cdh1* (*Cdh1*^*Gt wt*^) *or Cdh1-9A* (*Cdh1*^*9A*^), we introduced the P17/Cdh1 replacement vector together with pCAGGS-Cre (Araki et al., 1997) into *Cdh1*^+/GT^ ES cells using electroporation. The P17/Cdh1 replacement vector was designed to insert full length cDNA of *Cdh1*^*Gt wt*^ or *Cdh1*^*9A*^ between 5’ lox 66 and 3’ lox P. Point mutations that substituted Thr/Ser codons with Ala codons were introduced into *Cdh1* cDNA (Eurofins), generating the P17/Cdh1-9A replacement vector. After electroporation, ES cells were cultured in medium containing puromycin for 1 day to isolate cell lines that had undergone Cre-mediated recombination. Puromycin selection was performed twice at a 2-day interval. To detect the expression from the *Cdh1*^*wt*^ and *Cdh1*^*9A*^ knock-in (KI) alleles in the recombinant ES cell lines, we performed reverse transcription-PCR (RT-PCR) analysis using the primers 5NC-s2 (5’-TCGAACAGGCGCGGCGTGTT-3’) and mFzr as2 (5’-ATAGTCCTGGTCCATGGTG GAG-3’). The PCR product was cloned into the pGEM-T easy vector (Promega) and sequences were verified. Chimera mice were generated by morula injection (host ICR) of recombinant ES cells. Chimeric males were mated to C57BL/6N females and the progenies were genotyped by PCR using the following primers. Gs4 (5’-CCTCCACTACAGCAGCACG-3’) and Gas7 (5’-CTCCAAGGCCTTTGTGAGGC-3’) for the wild-type allele (0.4kb). Gs4 and SA6as (5 -CCGGCTAAAACTTGAGACCTTC-3’) for the knock-in allele (0.7kb).

### Preparation of testis extracts and immunoprecipitation

Testis extracts were prepared as described previously (Ishiguro et al., 2011). Briefly, testicular cells were suspended in low salt extraction buffer (20 mM Tris-HCl [pH 7.5], 100 mM KCl, 0.4 mM EDTA, 0.1% TritonX100, 10% glycerol, 1 mM β-mercaptoethanol) supplemented with Complete Protease Inhibitor (Roche). After homogenization, the soluble chromatin-unbound fraction was separated after centrifugation at 100,000*g* for 30 min. The solubilized chromatin fraction was collected after centrifugation at 100,000*g* for 30 min at 4°C.

The endogenous APC/C was immunoprecipitated from chromatin-unbound fraction of *Cdh1*^+/+^, *Cdh1*^Gt wt/Gt wt^, *Cdh1*^9A/9A^ mice testes, using 2 μg of mouse anti-CDC27 antibody and 50 μl of protein A-Dynabeads (Thermo-Fisher). The beads were washed with low salt extraction buffer. The bead-bound proteins were eluted with 40 μl of elution buffer (100 mM Glycine-HCl [pH 2.5], 150 mM NaCl), and then neutralized with 4 μl of 1 M Tris-HCl [pH 8.0].The immunoprecipitated proteins were run on 4-12 % NuPAGE (Thermo-Fisher) in MOPS-SDS buffer and immunoblotted. For the immunoblot of testes extracts, whole lysates were prepared in RIPA buffer and run on 8% Laemmli SDS-PAGE in Tris-Glycine-SDS buffer. Immunoblots were detected by secondary antibody VeriblotBlot for IP detection reagent (HRP) (ab131366, abcam, 1:3000 dilution), ECL rabbit IgG HRP-linked F(ab’)2 fragment, ECL mouse (NA9340, GE, 1:3000 dilution) or IgG HRP-linked F(ab’)2 fragment (NA9310, GE, 1:3000 dilution). Immunoblot image was developed using ECL prime (GE healthcare) and captured by FUSION Solo (VILBER).

### Antibodies

The following antibodies were used for immunoblot (IB) and immunofluorescence (IF) studies: rabbit anti-Actin (IB, 1:1000, Sigma A2066), mouse anti-CDC27 (IB, 1:1000, Abcam: ab10538), rabbit anti-Cdh1 (IB, IF, 1:1000, Abcam: ab118939), rabbit anti-CDC20 (IB, 1:1000, Bethyl:A301-180A), rabbit anti-SOX9 (IF, 1:1000, Milipore: Ab5535), rabbit anti-CyclinB1(IB, 1:1000, Santa Cruz: sc-595), mouse anti-PLK1(IB, 1:1000, Abcam ab17056), rabbit anti-SYCP1 (IF, 1:1000, Abcam ab15090), rabbit anti-γH2AX (IF, 1:1000, Abcam ab2893), rat anti-TRA98 (IF, 1:1000, ab82527), rabbit anti-CENPC (Ishiguro et al., 2011), rabbit anti-MEIKIN (IF, 1:1000) (Kim et al., 2015), rat anti-SYCP3 (Ishiguro et al., 2020), guinea pig anti-H1t (IF, 1:2000, kindly provided by Marry Ann Handel).

#### Histological Analysis

For, hematoxylin and eosin staining, testes, epididymis and ovaries were fixed in 10% formalin or Bouin solution, and embedded in paraffin. Sections were prepared on CREST-coated slides (Matsunami) at 6 μm thickness. The slides were dehydrated and stained with hematoxylin and eosin.

For Immunofluorescence staining, testes were embedded in Tissue-Tek O.C.T. compound (Sakura Finetek) and frozen. Cryosections were prepared on the CREST-coated slides (Matsunami) at 8 μm thickness, and then air-dried and fixed in 4% paraformaldehyde in PBS at pH◻7.4. The serial sections of frozen testes were fixed in 4% PFA for 5 min at room temperature and permeabilized in 0.1% TritonX100 in PBS for 10 min. The sections were blocked in 3% BSA/PBS, and incubated at room temperature with the primary antibodies in a blocking solution. After three washes in PBS, the sections were incubated for 1 h at room temperature with Alexa-dye-conjugated secondary antibodies (1:1000; Invitrogen) in a blocking solution. For paraffin embedded section, sections were deparaffinized, rinsed in distilled water and incubated with PBS for 30 minutes at 37◻. Then, sections were incubated in PBS containing for Proteinase K for 30 minutes at 37◻ and rinsed in distilled water. After treatement, immunofluorescence staining were performed with same procedure of frozen tissue sections.

TUNEL assay was performed using MEBSTAIN Apoptosis TUNEL Kit Direct (MBL 8445). DNA was counterstained with Vectashield mounting medium containing DAPI (Vector Laboratory). Lectin staining was done using Lectin from *Arachis hypogaea* FITC Conjugate (IF, 1:1000, Sigma: L7381).

### Immunostaining of spermatocytes

Spread nuclei from spermatocytes were prepared as described (Ishiguro et al., 2014). Briefly testicular cells were suspended in PBS, then dropped onto a slide glass together with an equal volume of 2% PFA, 0.2% (v/v) Triton X-100 in PBS, and incubated at room temperature in humidified chamber. The sides were then air-dried and washed with PBS containing 0.1% Triton-X100 or frozen for longer storage at −80ºC. The serial sections of frozen testes were fixed in 4% PFA for 5 min at room temperature and permeabilized in 0.1% TritonX100 in PBS for 10 min. The slides were blocked in 3% BSA/PBS, and incubated at room temperature with the primary antibodies in a blocking solution. After three washes in PBS, the sections were incubated for 1 h at room temperature with Alexa-dye-conjugated secondary antibodies (1:1000; Invitrogen) in a blocking solution. DNA was counterstained with Vectashield mounting medium containing DAPI (Vector Laboratory).

### Fluorescence *in situ* hybridization (FISH) on immunostained nuclei

For immuno-FISH, structurally preserved nuclei (SPN) from spermatocytes were prepared as described (Ishiguro et al., 2014) with modification. Briefly testicular cells were collected in PBS by mincing seminiferous tubules into small pieces with fine-tipped tweezers and then pipetting. After removal of tissue pieces, the cell suspension was filtered through a Cell strainer (BD Falcon) to remove debris. The cell suspension (~5 μl) was dropped onto a MAS-coated slide glass (Matsunami) and fixed with 10 μl of 2% Paraformaldehyde (PFA)/100 mM sucrose in PBS for 10 min followed by the addition of 1.5 μl of 1.25 M Glycine/PBS, and then air-dried at room temperature. Immediately before they were completely air-dried, the slide glasses were washed with PBS containing 0.1% Triton-X100 or frozen for longer storage at −80ºC.

Immuno-stained samples of SPN were fixed in 4% paraformaldehyde for 5 min, washed with PBS, and subjected to sequential dehydration through 70%, 80%, 90%, 100% ethanol. Immuno-stained SPNs were denatured in 50% formamide, 2× SSC at 72°C for 10 min. Hybridization was conducted with a fluorescence-labeled point probe in buffer containing 50% formamide, 2× SSC, 20% dextran sulfate at 37°C for 12-16 h. The slides were washed sequentially at room temperature in 2× SSC for 1 min, 0.4× SSC/0.3% Tween20 solution for 2 min, and 2×SSC at room temperature for 1 min. The mouse point probe derived from BAC clone RP23-6I6 detects the mid region of chromosome 3.

### Imaging

Immunostaining images were captured with DeltaVision (GE Healthcare). The projection of the images were processed with the SoftWorx software program (GE Healthcare). For Fig S2B, images were captured with Zeiss LSM-710 confocal microscope and processed with ZEN software. All images shown were Z-stacked. Bright field images were captured with OLYMPUS BX53 fluorescence microscope and processed with CellSens standard program. For counting seminiferous tubules, immunostaining images were captured with BIOREVO BZ-X710(KEYENCE), and processed with BZ-H3A program.

---

**Supplementary Figure 1.**
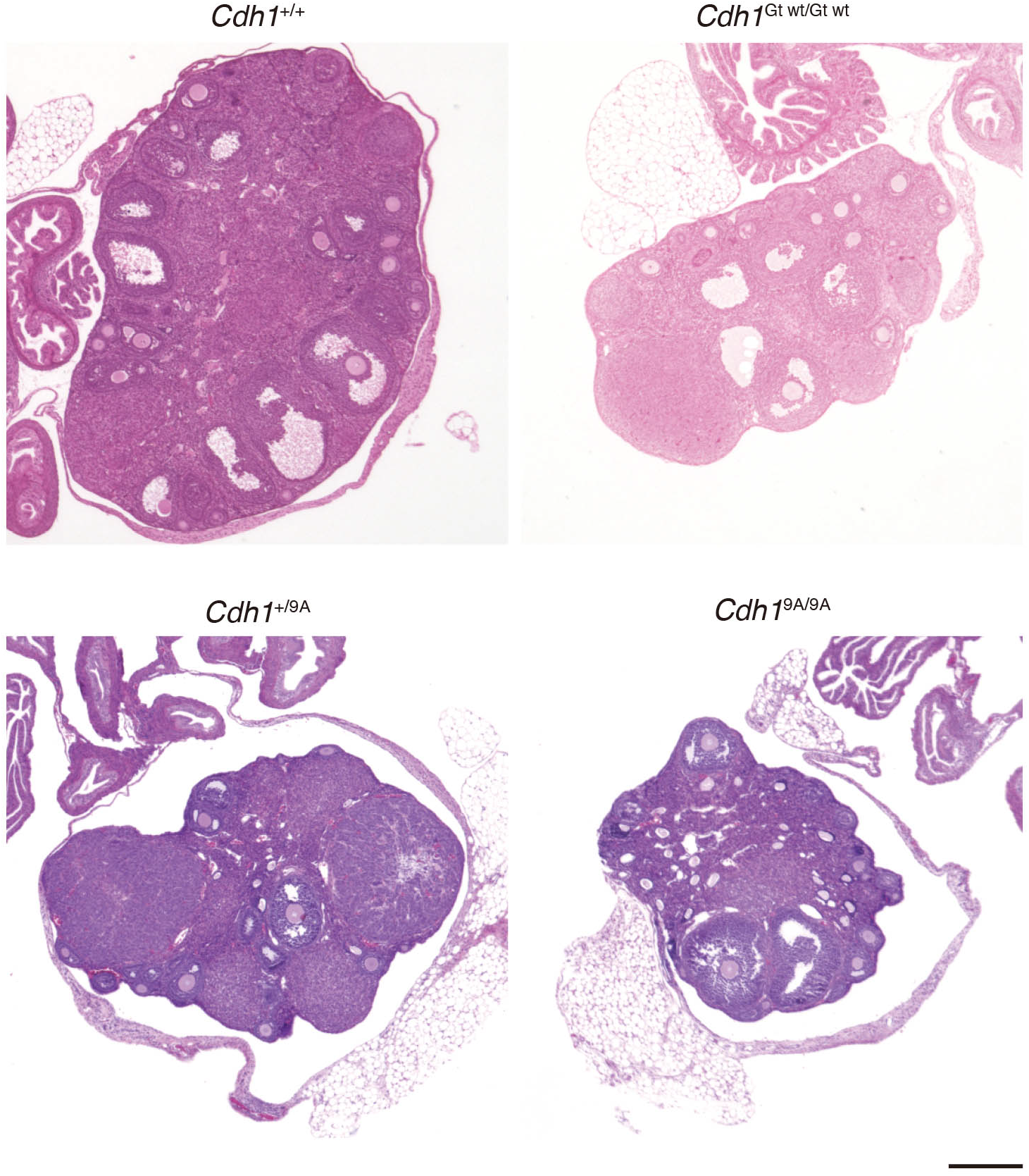
*Cdh1*^9A/9A^ KI females showed no overt defects in adult ovaries. Hematoxylin and Eosin stained sections of WT, *Cdh1*^Gt wt/Gt wt^, *Cdh1*^+/9A^ and *Cdh1*^9A/9A^ KI ovaries (8-weeks). Scale bar: 300μm

**Supplementary Figure 2.**
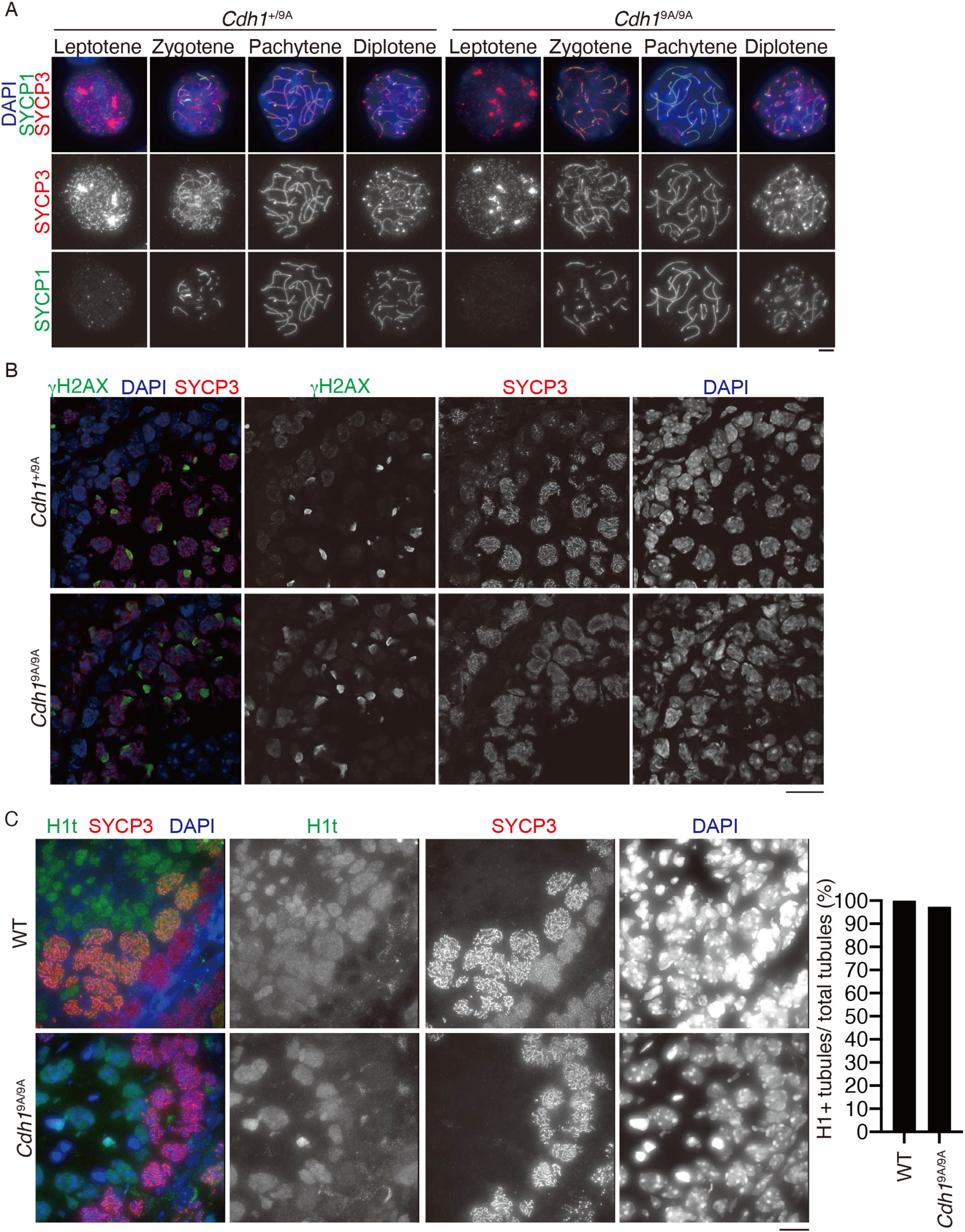
Normal progression of meiotic prophase in *Cdh1*^9A/9A^ KI males. **(A)** Spread nuclei from the testes of WT and *Cdh1*^9A/9A^ KI mice were immunostained for SYCP3, SYCP1 and DAPI. Scale bar: 5 μm. **(B)** Seminiferous tubule sections of *Cdh1*^+/9A^ and *Cdh1*^9A/9A^ KI mice (3-week-old) were immunostained for SYCP3, γH2AX and DAPI. Scale bar: 20 μm. **(C)** Seminiferous tubule sections in WT and *Cdh1*^9A/9A^ KI mice (5-week-old) were immunostained for SYCP3, Histone H1t, SYCP1 and DAPI. Quantification of the seminiferous tubules that have H1t +/SYCP3+ cells per the seminiferous tubules that have SYCP3+ spermatocyte cells in WT (8w: n= 1) and *Cdh1*^9A/9A^ KI mice (8w: n= 1) testes (bar graph). Scale bar: 15 μm.

**Supplementary Figure 3.**
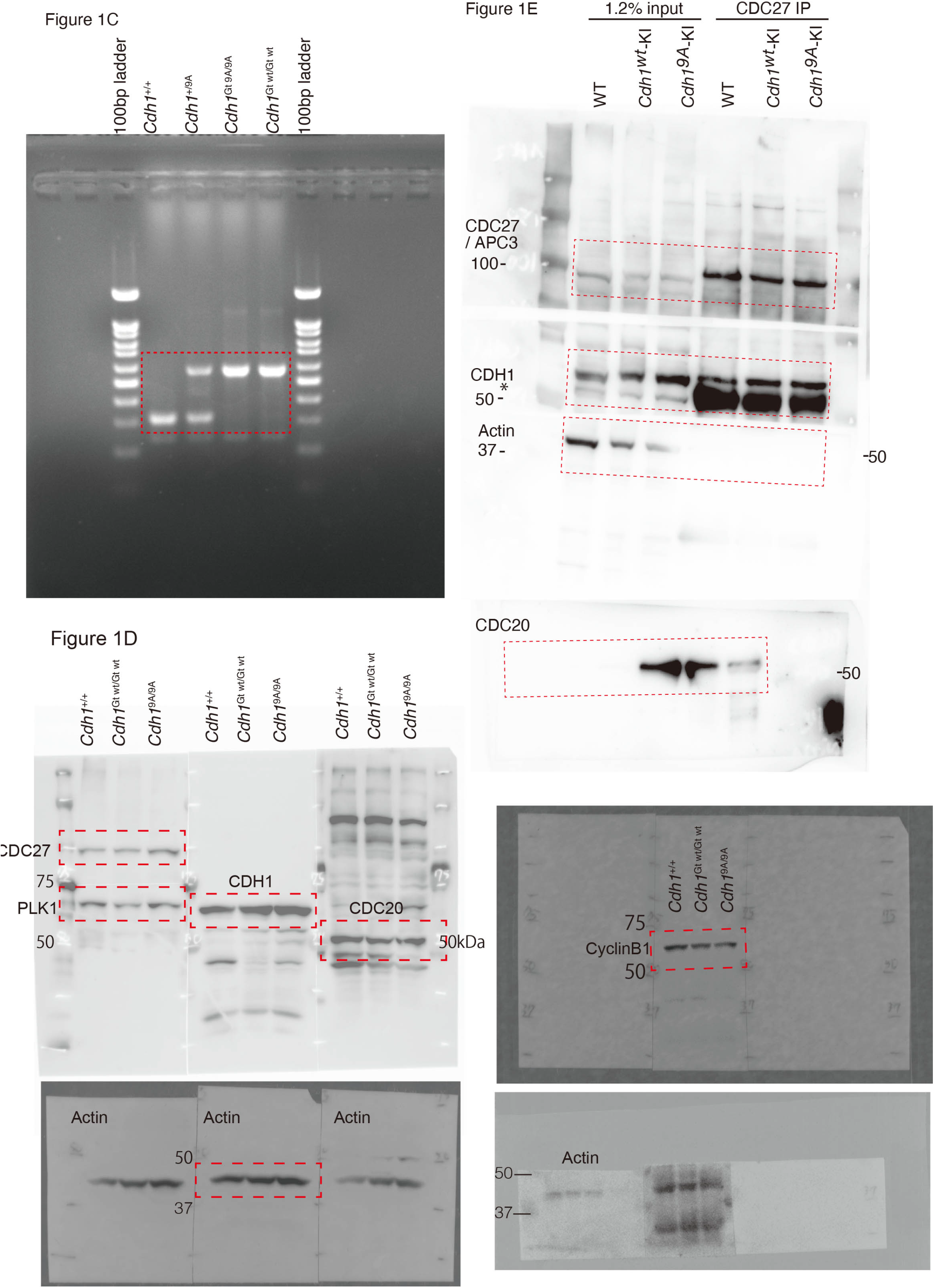
Full-length / uncropped images. Full-length / uncropped images of agarose gel (Fig1C) and immunoblots (Fig1D, E) are shown.

